# Disruption of relapse to alcohol seeking by aversive counterconditioning following memory retrieval

**DOI:** 10.1101/841536

**Authors:** Koral Goltseker, Hen Handrus, Segev Barak

## Abstract

Relapse to alcohol abuse is often caused by exposure to potent alcohol-associated cues. Therefore, disruption of the cue-alcohol memory can prevent relapse. It is believed that memories destabilize and become prone for updating upon their reactivation through retrieval, and then re-stabilize within 6 h during a “reconsolidation” process. We recently showed that relapse to cocaine seeking could be prevented by counterconditioning the cocaine-cues with aversive outcomes following cocaine-memory retrieval, in a place conditioning paradigm. However, to better model addiction-related behaviors, self-administration models are necessary. Here, we demonstrate that relapse to alcohol seeking can be prevented by aversive counterconditioning conducted during alcohol-memory reconsolidation, in conditioned place preference (CPP) and operant self-administration paradigms, in mice and rats, respectively. We found that the reinstatement of alcohol-CPP was abolished only when aversive counterconditioning with water-flooding was given shortly after alcohol-memory retrieval. Furthermore, rats trained to lever-press for alcohol showed decreased context-induced renewal of alcohol-seeking responding when the lever-pressing was counterconditioned with foot-shocks, shortly, but not 6 h, after memory retrieval. These results 0suggest that aversive counterconditioning can prevent relapse to alcohol seeking only when performed during alcohol-memory reconsolidation, presumably by updating, or replacing, the alcohol memory with aversive information. Also, we found that aversive counterconditioning preceded by alcohol-memory retrieval was characterized by upregulation of brain-derived neurotrophic factor (*Bdnf)* mRNA expression in the medial prefrontal cortex, suggesting that *Bdnf* plays a role in the memory updating process.

## Introduction

Alcohol addiction is a relapsing disorder; even with successful treatments, nearly 70% of patients relapse within the first year of abstinence, with no available effective preventive treatment (Marlatt, 1996; Bottlender and Soyka, 2004; Hendershot et al., 2011; Sinha, 2011; Witteman et al., 2015; Batra et al., 2016). Relapse to alcohol abuse is often caused by exposure to the environmental cues that became associated with the reinforcing effects of alcohol (Miller and Gold, 1998; Hyman et al., 2006; Loeber et al., 2006; Witteman et al., 2015). Therefore, disruption of the cue-alcohol association is expected to reduce or even prevent relapse.

Aversion therapy is one way to attenuate cue-induced drug/alcohol craving and relapse (Cannon and Baker, 1981; Elkins, 1991; Howard et al., 1991; Elkins et al., 2017). In this approach, a cue previously associated with the reinforcing effects of alcohol, is re-associated (counterconditioned) with an aversive outcome, such as a mild electrical shock or induction of nausea (Wiens et al., 1976; Miller, 1978; Cannon et al., 1981). Counterconditioning was shown to be more potent than extinction in suppressing relapse in animal models and humans (Van Gucht et al., 2010; Tunstall et al., 2012), and to help alcoholic patients to stay sober for a long period of time (Elkins et al., 2017). However, the suppressive effect is temporary (Bouton and Peck, 1992; Brooks et al., 1995; van Dis et al., 2019). Suggestively, aversive counterconditioning leads to formation of a new cue-aversion association that competes with the cue-alcohol association for behavioral expression (Bouton and Peck, 1992; Brooks et al., 1995). Thus, the persistent cue–alcohol memory can recover, reinstating cue-induced craving and relapse.

As an alternative, efficient suppression of unwanted memories can be possibly achieved by updating the original alcohol memory itself (Lee et al., 2017). Current concepts suggest that memories undergo de-stabilization upon their reactivation via memory retrieval, and then re-stabilize within a few hours in a process termed “reconsolidation” (Przybyslawski and Sara, 1997; Nader et al., 2000). While in a temporary active state (termed “reconsolidation window”), memories can be disrupted, updated, or strengthened (Nader and Hardt, 2009; Nader, 2015; Dunbar and Taylor, 2017b; Lee et al., 2017; Gisquet-Verrier and Riccio, 2018), providing a promising therapeutic approach for relapse prevention in addiction (Tronson and Taylor, 2013; Vousden and Milton, 2017; Exton-McGuinness and Milton, 2018). It is established that memory reconsolidation can be disrupted by pharmacological manipulations (Nader et al., 2000; Dunbar and Taylor, 2017b). However, the toxicity or side effects of most relevant pharmacological amnesic agents (mainly protein synthesis inhibitors) (Nader et al., 2000; Dunbar and Taylor, 2017b), facilitated the development of behavioral manipulations to interfere with memory reconsolidation (Ma et al., 2012; Xue et al., 2012; Millan et al., 2013; Sartor and Aston-Jones, 2014; Das et al., 2015; Luo et al., 2015; Cofresi et al., 2017; Goltseker et al., 2017; Das et al., 2018; Gera et al., 2019).

We recently showed that applying aversive counterconditioning following the retrieval of cocaine-memories prevented reinstatement of cocaine seeking (Goltseker et al., 2017), suggesting that a cue-aversion memory replaced the cue-drug memory trace, rather than formed a competing memory. We also showed that cocaine seeking was reinstated when counterconditioning was given without, 5-h after, or before the reactivation of cocaine-memories, indicating that counterconditioning should be applied during the ‘reconsolidation window’ to prevent relapse. These findings were also in line with the evidence from a study, in which re-association of the alcohol cues with aversion upon alcohol memory retrieval yielded reduction in craving for alcohol in hazardous drinkers (Das et al., 2015; Das et al., 2018).

In our previous cocaine study (Goltseker et al., 2017) we used place conditioning, in which the drug is injected to, rather than consumed voluntary by the animal. Here, we sought to prevent relapse to alcohol seeking by incorporating aversive information into the cue-alcohol memory, in both place conditioning, and operant self-administration paradigms, with the latter serving as a better model for addiction-related behaviors (Panlilio and Goldberg, 2007; Goltseker et al., 2019). Specifically, we tested whether aversive counterconditioning of the alcohol-associated cues/context, conducted shortly after the retrieval of alcohol memories, can prevent reinstatement of alcohol-conditioned place preference (CPP) in mice, or renewal of alcohol seeking in an operant self-administration paradigm in rats. We also investigated the molecular correlates of the memory replacement mechanism, by assessing the expression of brain-derived-neurotrophic factor (*Bdnf*) and zinc-finger protein 268 (*Zif268*), genes previously implicated in memory processing (Lee et al., 2004; Lee et al., 2005; Tedesco et al., 2014; Radiske et al., 2015; Radiske et al., 2017), in the brain regions associated with memory reconsolidation.

## Materials and Methods

### Animals

Male and female C57BL/6 mice (25-30 g), housed 3-4/cage, and male and female Wistar rats (250-450 g), housed 2/cage, were bred at Tel-Aviv University Animal Facility (Israel), and kept under a 12-h light-dark cycle (lights on at 7 a.m.) with food and water available ad libitum. All experimental protocols were approved by, and conformed to, the guidelines of the Institutional Animal Care and Use Committee of Tel Aviv University, and to the guidelines of the NIH (animal welfare assurance number A5010-01). All efforts were made to minimize the number of animals used.

### Drugs and reagents

Ethanol Absolute (Gadot, Israel) was dilluted to 20% v/v solution in tap water (for voluntary intake) or in sterile 0.9 M NaCl saline (for systemic injections). In CPP experiments, a dose of 1.8 g/kg alcohol was administered intraperitoneally in a volume of 10 ml/kg. This dose was previously shown to produce alcohol-induced CPP (Neasta et al., 2010; Morisot et al., 2018). An equivalent volume of saline was used for control administrations. Fast SYBR Green Master Mix, TRIzol reagent, and RevertAid kit were supplied by Thermo-Fisher Scientific. DNA oligonucleotides (qRT-PCR primers) were obtained from Sigma-Aldrich.

### Apparatus

#### Place conditioning

All place conditioning experiments were performed in open-ceiling Plexiglas boxes (30 × 30 × 20 cm) divided non-hermetically into two equal-sized compartments by a sliding door. The compartments differed from each other by the wall pattern (horizontal vs. vertical b/w stripes; 1 cm width) and the floor surface (white textured plastic; with bulging circles vs. bulging stripes). The horizontal stripes pattern on the walls was always matched with the bulging circles on the floor, whereas the vertical stripes – with the bulging stripes. During the water-flood trials, the ceiling was covered with a removable transparent Plexiglas sheet to prevent mice from escaping. Each Plexiglas box was placed in a sound-attenuating chamber equipped with a LED light stripe on the walls and a ceiling camera that registered mouse behavior. Data were recorded by Ethovision XT 11.5 (Noldus, Wageningen, Netherlands).

#### Operant self-administration

All instrumental self-administration experiments were performed in operant conditioning modular chambers for rats (Med Associates Inc., Georgia, VT), equipped with two levers, a liquid dispenser and a receptacle, Plexiglas walls, grid floor, and a waste pane, and placed in sound-attenuating cubicles with a LED light stripe on the walls. The unadjusted manufacturer setting with the LED-lights turned off served as Context A. For Context B, the Plexiglas walls were covered with pattern sheets (vertical b/w stripes; 1 cm width), the LED-lights turned on, the waste pan removed, and a carton sheet sprinkled with eucalyptus oil was inserted under the grid floor. During the counterconditioning phase, the grid floor was connected to aversive stimulator/scrambler modules (ENV-414; Med Associates Inc., Georgia, VT). Data were recorded by Med-PC IV Software Suite (Med Associates Inc., Georgia, VT).

### Procedures

#### Aversive counterconditioning in a place conditioning paradigm

All mice were habituated to daily i.p. saline administrations 5 days prior to the beginning of the procedure.

##### Baseline test (day 1)

On the first day, the sliding door was retracted and mice were allowed to explore the entire apparatus freely for 30 min. Animals that spent >70% of time in either of the compartments were excluded from the study (2 mice from Experiment 1). This allowed the use of an unbiased design, in which the 2 compartments are: a) equally preferred before conditioning as indicated by the group average (unbiased apparatus); b) pseudo-randomly assigned to the experimental conditions (unbiased assignment procedure) (Cunningham et al., 2003). Each place conditioning chamber was assigned to one sex.

##### Alcohol place conditioning (days 2–9)

Training started 24 h after the Baseline test with one session per day over 8 days, with the sliding door closed. On days 3, 5, 7, and 9, mice were administered with alcohol (1.8 g/kg; i.p.) and immediately confined to the paired compartment for 5 min. On the alternate days (days 2, 4, 6, and 8), mice were administered with saline and were confined to the unpaired compartment for the same duration as on the alcohol-conditioning day. Paired compartments were counterbalanced.

##### Place preference test 1 (day 10)

Place preference test was identical to the Baseline stage, and served to index alcohol-CPP (Cunningham et al., 2003). Preference was defined as an increase in the percent of time spent in the alcohol-paired compartment during the Place preference test 1 compared to the Baseline test. Mice that did not show CPP (minimum of 5% change) were excluded from the experiment (3 from Experiment 1; 4 from Experiment 2).

##### Memory retrieval and aversive counterconditioning (days 11-14)

Prior to this stage, mice were assigned to different experimental conditions, Retrieval and No Retrieval (matched for CPP scores and sex). Training began 24 h after Place Preference Test 1, with a session per day over 4 days. Specifically, on days 12 and 14, mice of the Retrieval group were confined to the alcohol-paired compartment for 3 min, whereas mice of the control, No Retrieval group were handled. Mice were then returned to their home cages and held in a service room adjoined to the experiment room. Counterconditioning started 45 min after the memory retrieval. Mice were placed in the alcohol-paired compartment, and shortly after the session started, two liters of water (18-20°C) were poured into the opposite (unpaired) compartment. As a result, water flowed under the sliding door and gradually flooded the paired compartment (where the mouse was located) to ∼2 cm height (for the detailed description of the procedure see Goltseker and Barak, 2018). Mice from the Retrieval group remained in the compartment for 12 min, whereas mice from the No Retrieval group - for 15 min, so that all mice spent a total of 15 min in the alcohol-associated compartment on the paired days. On days 11 and 13, mice were placed in the unpaired compartment, with no water poured into the conditioning chamber.

##### Place preference test 2 (day 15)

This stage was identical to the previous test stages, and confirmed that counterconditioning led to the loss of alcohol-CPP.

##### Reinstatement test (day 16)

On this day, all mice received a prime administration, half of the conditioning dose (Mueller and Stewart, 2000; Shoblock et al., 2011) of alcohol (0.9 g/kg) immediately before a 30-min test session with free access to both compartments. Reinstatement of alcohol-CPP is considered to model relapse to alcohol seeking (Katner and Weiss, 1999; Roger-Sanchez et al., 2012; Ben Hamida et al., 2018). Reinstatement was defined as an increase in the percentage of time spent in the alcohol-paired compartment during the Reinstatement test, as compared to the Place preference test 2.

#### Aversive counterconditioning in an alcohol operant self-administration paradigm

##### Acquisition of operant alcohol self-administration

This procedure was previously described by Simms et al. (2010) (also see Barak et al., 2013; Carnicella et al., 2014). Briefly, during the first 7 weeks, rats were given intermittent access to 20% alcohol in the 2-bottle choice paradigm. Operant alcohol self-administration training in Context A started with 6 14-h overnight sessions on a fixed ratio 1 (FR1) schedule of reinforcement (0.1 ml 20% alcohol solution (v/v) following a single active lever press), given every other day over two weeks. Rats were then trained in daily sessions given 5 days a week, first in 60-min FR1 sessions for 3 weeks (15 sessions), then in 60-min FR3 sessions (alcohol delivery following 3 active lever presses) for 2 weeks (10 sessions), and then in daily 30-min FR3 sessions for another 2 weeks (10 sessions). The number of presses on each lever (active and inactive), as well as the number of reward deliveries, were recorded. Rats that attained less than 10 alcohol deliveries in average during the last 6 training sessions were excluded from the experiment (2 from Experiment 3; 1 from Experiment 4) (Carnicella et al., 2014). Each operant chamber was assigned to one sex.

##### Memory retrieval and aversive counterconditioning

Before this stage, rats were given 10 days of abstinence from alcohol intake, and were kept undisturbed in the animal facility (Barak et al., 2013). Then, rats were assigned to the experimental conditions, Retrieval and No Retrieval (matched for the number of active lever presses and sex). For the Retrieval group, an empty bottle with 0.2-ml of 20% alcohol applied on its tip was given in the home cage for 10 min. Rats of the No Retrieval group received only a bottle containing water. This procedure was previously shown to initiate the reconsolidation of alcohol-associated memories (Barak et al., 2013). Forty-five minutes later (Experiment 3), or 6 h later (Experiment 4), rats from both groups were placed in the Context B operant chambers. At the beginning of each session, rats were allowed to consume a non–pharmacologically active alcohol prime (0.2 ml of 20% alcohol) from the receptacle, in order to facilitate operant response on the levers presented into the chamber. Similar to the final acquisition sessions, counterconditioning was performed under an FR3 reinforcement schedule. However, here, each third press on the active lever resulted in the delivery of a mild foot-shock (0.25 mA; 0.5 sec), rather than alcohol reward. The maximum number of foot-shocks per session was limited to six, to avoid high variability between rats. Upon meeting the criterion, or 30 min after the beginning of the session, rats were returned to the home cages. Lever presses on the inactive lever were not reinforced. Four counterconditioning sessions with or without memory retrieval were given over 4 days.

##### Reinstatement tests

Relapse to alcohol seeking was assessed in a reinstatement test stage, conducted in Context A or in Context B, 24 or 48 h after the last counterconditioning session. Rats were placed in the operant chambers for a 30-min session, similarly to the self-administration acquisition sessions, except that no alcohol was delivered after either lever was pressed. In addition, an alcohol prime (0.2 ml, 20%) was non-contingently presented at the beginning of the session, as a -odor-taste cue.

#### Quantitative reverse transcriptase polymerase chain reaction (qRT-PCR)

Following brain dissection, tissue samples were immediately snap-frozen in liquid nitrogen and stored at −80°C until use. Frozen tissues were mechanically homogenized in TRIzol reagent and total RNA was isolated from each sample according to the manufacturer’s recommended protocol. mRNA was reverse transcribed to cDNA using the Reverse Transcription System and RevertAid kit. Plates of 96 wells were prepared for the SYBR Green cDNA analysis using Fast SYBR Master Mix. Samples were analyzed in triplicate/duplicate with a Real-Time PCR System (StepOnePlus; Applied Biosystems), and quantified against an internal control gene, *Gapdh*. We used the following reaction primers sequences: *Egr1*(*Zif268*) forward, 5′-TGAGCACCTGACCACAGAGTC-3′; reverse, 5′-TAACTCGTCTCCACCATCGC-3′; *Bdnf IV* forward, 5′-GCAGCTGCCTTGATGTTTAC-3′; reverse, 5′-CCGTGGACGTTTACTTCTTTC-3′; *Gapdh* forward, 5′-CCAGAACATCATCCCTGC-3′; reverse, 5′-GGAAGGCCATGCCAGTGAGC-3′. Thermal cycling was initiated with incubation at 95°C for 20 s (for SYBR Green activation), followed by 40 cycles of PCR with the following conditions: heating at 95°C for 3 s and then 30 s at 60°C. Relative quantification was calculated using the ΔΔCt method.

#### Experimental design and data analysis

##### Experiment 1

Following a Baseline test, mice were trained for alcohol-CPP in 4 conditioning sessions. The expression of alcohol-CPP was subsequently confirmed in Place preference test 1. Next, mice were briefly re-exposed to the alcohol-paired compartment (Retrieval group) or handled (No Retrieval group), followed 45 min later by water-flood aversive counterconditioning. The retrieval-counterconditioning sessions were repeated twice. The suppression of alcohol-CPP was confirmed in Place preference test 2. Next, relapse to alcohol seeking was assessed in an alcohol-primed Reinstatement test. Place preference was indexed by percentage of time spent in the alcohol-paired compartment (time spent in the alcohol-paired compartment X 100 / total test time). Expression of alcohol-CPP following conditioning was analyzed by a paired t-test, comparing the place preference during the Baseline (pre-conditioning) and Place preference test 1 (post-conditioning). Place preference in the tests conducted after aversive counterconditioning was analyzed by a mixed-model ANOVA, with a between-subject factor of Group (Retrieval, No Retrieval) and a repeated measures factor of Test (Place preference test 2, Reinstatement test). Significant ANOVA was followed by a Student–Newman–Keuls (SNK) post-hoc test.

##### Experiment 2

This experiment was identical to Experiment 1, except that mice were euthanized shortly after aversive counterconditioning, preceded or not by alcohol-memory retrieval. The dorsal hippocampus and medial prefrontal cortex (mPFC) were collected and processed for RT-qPCR. Target gene mRNA levels were normalized to *Gapdh* (He and Ron, 2006; Barak et al., 2015; Even-Chen et al., 2017; Ziv et al., 2019), followed by normalization to the control condition (No Retrieval+No Counterconditioning). Levels of *Bdnf* and *Zif268* mRNA expression per each brain region were analyzed using a factorial MANOVA, with between-subjects’ factors of Group (Retrieval, No Retrieval) and Training (Counterconditioning, No Counterconditioning). A significant MANOVA was followed by an SNK post-hoc test per gene. Data from one mouse were excluded from the analysis due to RNA degradation.

##### Experiment 3

Following 7-weeks of alcohol consumption in a home cage, rats were trained to lever press for alcohol in an operant alcohol self-administration procedure in Context A for 9 weeks. After 10 days of abstinence, rats were briefly re-exposed or not to the alcohol -odor-taste cues in the home cage (Barak et al., 2013). Forty-five minutes later, rats were placed in Context B, in which a lever press triggered delivery of a mild foot-shock. After 4 aversive counterconditioning sessions (a session per day), alcohol-seeking was assessed in Context A and Context B. Active and inactive lever presses during the tests were analyzed by a mixed-model ANOVA, with a between-subject factor of Group (Retrieval, No Retrieval) and a repeated measures factor of Test (Test A, Test B). A significant ANOVA was followed by an SNK test. The number of active and inactive lever presses during the last 6 alcohol-training sessions was analyzed with a mixed-model ANOVA, with a between-subjects factor of Lever (Active, Inactive) and a repeated measures factor of Day (Days 1-6). The number of active lever-presses and foot-shocks during the counterconditioning stage were analyzed by a mixed-model ANOVA, with a between-subject factor of Group (Retrieval, No Retrieval) and a repeated measures factor of Day (Days 1-4).

##### Experiment 4

This experiment was identical to Experiment 3, except that the counterconditioning sessions were conducted 6 h after the memory retrieval sessions.

Sex distributed approximately equally across the experiments, and was initially analyzed as a factor; however, all analyzes did not yield a main effect of sex or any interaction with other factors (p’s >0.05). Therefore, data were collapsed across this factor.

## Results

### Experiment 1: Aversive counterconditioning within the “reconsolidation window” prevents relapse to alcohol seeking in a place conditioning paradigm

First, we tested whether aversive counterconditioning performed shortly after the retrieval of alcohol-associated memories could prevent the reinstatement of alcohol seeking, in a place conditioning paradigm. This behavioral paradigm has been widely used to study the reinforcing properties of drugs and alcohol (van der Kooy et al., 1983; Cunningham et al., 2003; Barak et al., 2011; Gibb et al., 2011), and to explore mechanisms of relapse to alcohol seeking and its prevention (Shoblock et al., 2011; Napier et al., 2013; Ben Hamida et al., 2018). The place conditioning paradigm can be used to induce both appetitive conditioned place preference (CPP), or aversive conditioned place aversion (CPA) (Cunningham et al., 2006; Tzschentke, 2007). Recently, we developed a 2-phase procedure, in which CPP was induced by cocaine, and was then counteracted in a lithium chloride (LiCl) CPA procedure (Goltseker et al., 2017).

Here, we applied a non-pharmacological aversive stimulus (water flood (Goltseker and Barak, 2018)) as an alternative to LiCl, since the pharmacological effects of the latter have been implicated in synaptic plasticity (Basselin et al., 2006), memory formation (Dewachter et al., 2009), memory reconsolidation (Wu et al., 2011), and memory extinction (Goltseker et al., 2017).

As presented in Figure 1A (experimental timeline), following a baseline test, all mice were first trained in an alcohol-CPP procedure, and then tested in Place preference test 1. As expected, mice displayed alcohol-CPP, as reflected by an increased percentage of time spent in the alcohol-paired compartment during Place preference test 1, compared to baseline (Figure 1B; t(19)=-8.14; p<0.00001). In the second phase, mice underwent place counterconditioning with water-flood, 45 min after memory retrieval by a brief re-exposure to the alcohol-paired compartment (Retrieval group) or handling (control, No Retrieval group). The 45-min interval was chosen for the counterconditioning to be held within the ‘reconsolidation window’ (Nader and Hardt, 2009; Xue et al., 2012; Lee et al., 2017). The subsequent Place preference test 2 confirmed that aversive counterconditioning led to the loss of place preference in both groups (Figure 1B).

**Figure 1.**
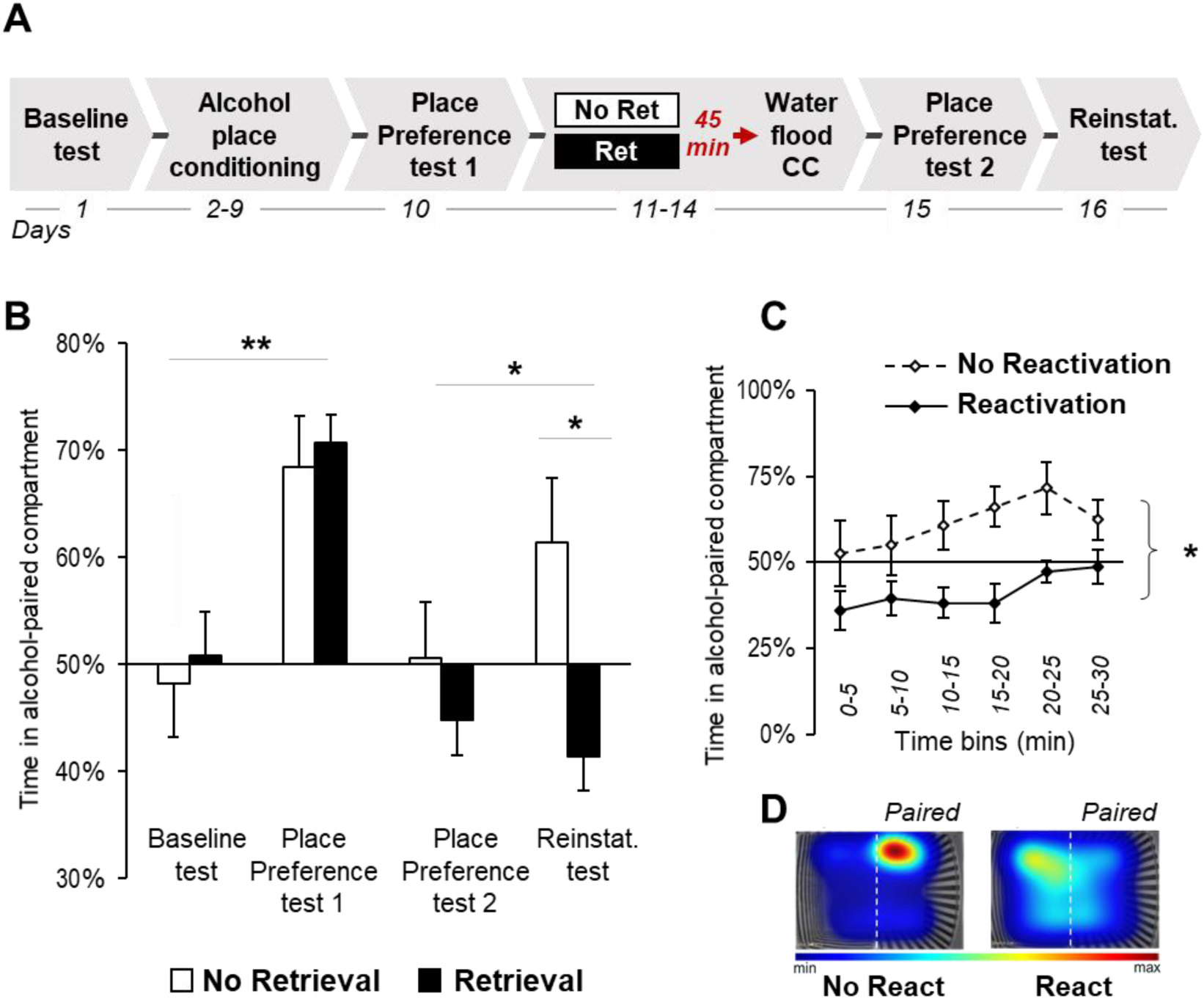
Aversive counterconditioning conducted shortly after retrieval of alcohol-associated memories prevents reinstatement of alcohol-conditioned place preference in mice. **A.** Schematic representation of the experimental procedure (CC, counterconditioning; Ret, retrieval). Mice were first trained for alcohol-CPP, and then received water-flood CPA training, conducted 45 min after a memory retrieval session consisting of a 3-min re-exposure to the alcohol-associated compartment (Retrieval group), or handling (control, No Retrieval group). Place preference tests were conducted after completion of the place conditioning and counterconditioning stages. Reinstatement of alcohol-CPP was assessed in a prime-induced test 24 h after validation of CPP loss. **B-C.** Place preference/aversion scores, expressed as mean * S.E.M. of the percent of time spent in the alcohol/water-paired compartment; *p<0.05; **<0.01; n=10 per group. **B.** Place preference/aversion scores during the entire tests sessions (30 min). **C.** Place preference/aversion scores during the Reinstatement test at 5-min temporal resolution. **D.** Representative heat maps depict the location of mice from No Retrieval vs Retrieval groups during the Reinstatement test. Heat map scale bar represents the normalized time spent at each XY coordinate during the tests.

Next, we tested whether alcohol-CPP would be reinstated by a prime administration of alcohol (0.9 g/kg), an established measure of relapse to alcohol seeking (Shoblock et al., 2011; Venniro et al., 2016; Ben Hamida et al., 2018). We found that while the No Retrieval group showed reinstatement of alcohol-CPP, the Retrieval group did not reinstate the preference to the alcohol-paired compartment, and spent significantly less time in the alcohol-paired compartment compared to the No Retrieval group (Figure 1B; Place preference test 2 vs. Reinstatement test: a mixed model ANOVA, a main effect of Group [F(1,18)=5.49, p<0.05], and a Group X Test interaction [F(1,18)=4.42, p<0.05], but no main effect of Test [F(1,18)=1.16, p>0.05]. Post-hoc analysis: No Retrieval group [Place preference test 2 vs Reinstatement test, p<0.05]; No Retrieval vs Retrieval group [Reinstatement test, p<0.05]).

Further analysis of the mice performance during the Reinstatement test per 5 min time bins, confirmed that the Retrieval group spent less time in the alcohol-paired compartment compared to the No Retrieval group during the 30-min test session (Figure 1C; a mixed-model ANOVA, main effects of Group [F(1,18)=8.91, p<0.01], and Time [F(5,90)=2.84, p<0.05], but no Group X Time interaction [F(5,90)=0.75, p>0.05]). Together, these results indicate that aversive counterconditioning prevents reinstatement of alcohol seeking, only when applied after the retrieval of alcohol-associated memories.

### Experiment 2: Aversive counterconditioning following alcohol-memories retrieval increases *Bdnf* expression in the mPFC

Next, we assessed the mRNA expression of *Zif268* and *Bdnf*, genes implicated in memory processing (Lee et al., 2004; Lee et al., 2005; Tedesco et al., 2014; Radiske et al., 2015; Radiske et al., 2017). We chose to focus on the mPFC and dorsal hippocampus, known to be involved in memory reconsolidation (Bozon et al., 2003; Lee et al., 2004; Tedesco et al., 2014; Radiske et al., 2015; Radiske et al., 2017), and in formation and expression of CPP or CPA (Zavala et al., 2003; Zhou and Zhu, 2006; Zarrindast et al., 2010; Chen et al., 2013; Otis et al., 2014; Slaker et al., 2015). Mice were trained in the alcohol-CPP procedure (Figure 2A; experimental design), and showed preference for the alcohol-paired compartment (Baseline test vs Place preference test 1 [t(22)=9.66, p<0.00001]). On the next day, one group of the animals was re-exposed to the alcohol-paired compartment (Retrieval group), whereas another group was briefly handled (No Retrieval group). Next, mice from each group were either subjected to a water-flood aversive counterconditioning session, or handled as a no-counterconditioning control. Thirty minutes after the end of the counterconditioning session (i.e., 90 min after memory retrieval) all mice were euthanized, and the mPFC and dorsal hippocampus were collected for further analysis of *Bdnf* and *Zif268* mRNAs expression.

**Figure 2.**
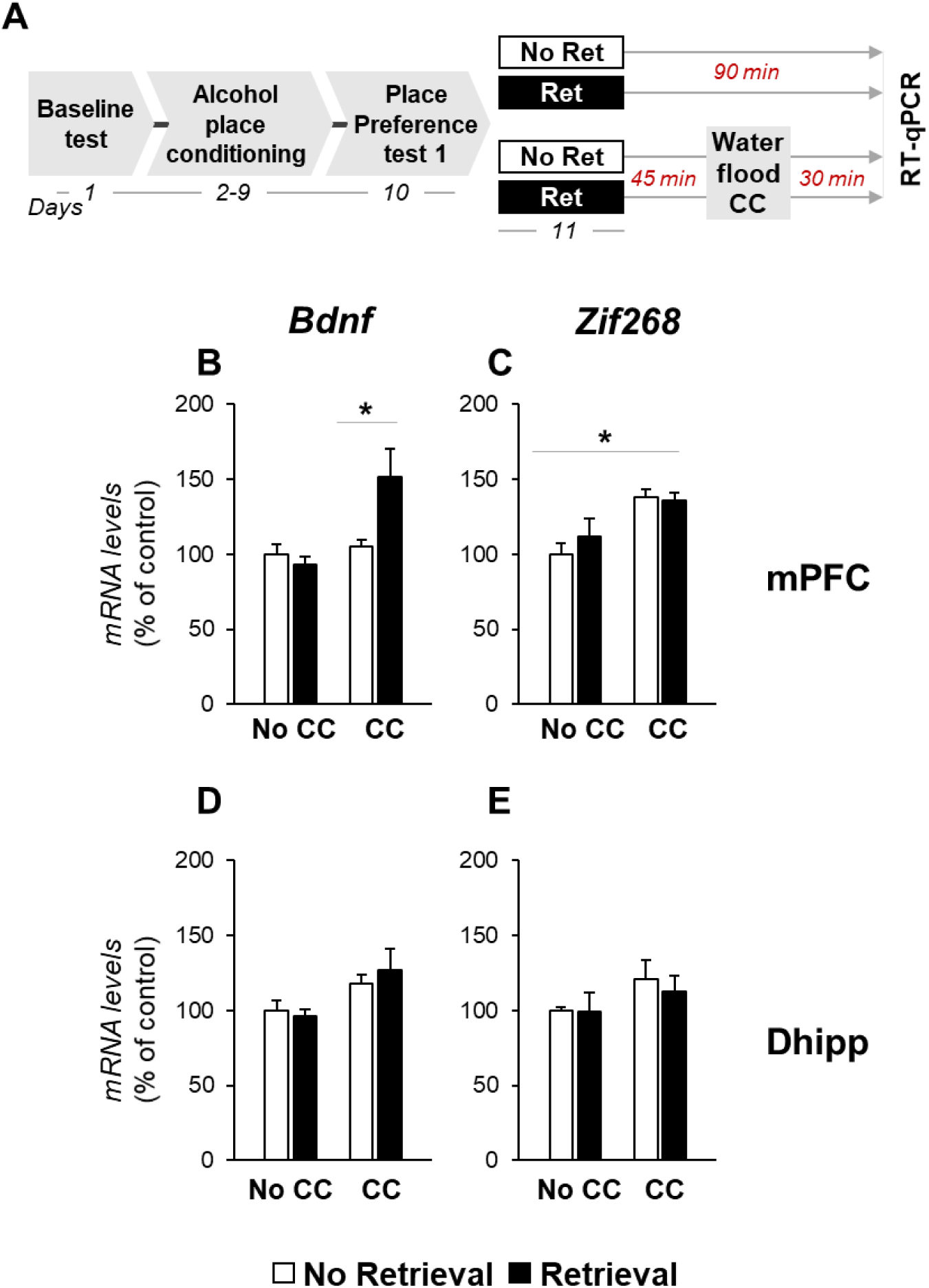
Aversive counterconditioning conducted shortly after retrieval of alcohol-places memories upregulates *Bdnf* mRNA expression in the mPFC in mice. **A.** Schematic representation of the experimental procedure (CC, counterconditioning; Ret, retrieval). Mice were first trained for alcohol-CPP, and then did or did not recieve place counterconditioning with water-flood, conducted 45 min after memory retrieval (Retrieval group), or handling (No Retrieval group). Ninety minutes after memory retrieval, brain tissues were collected for RT-qPCR analysis. **B-E.** mRNA levels of *Bdnf* and *Zif268* in the mPFC (B-C) and the dorsal hippocampus (D-E), were normalized to *Gapdh*, and are expressed as mean + S.E.M. of the percent of change from the control group (No Retrieval-No CC); *p<0.05; n=5-6 per group.

We found that in the mPFC, the increase in the expression of *Bdnf* following counterconditioning was dependent on prior memory retrieval (Figure 2B), whereas the increased levels of *Zif268* following counterconditioning were retrieval-independent (Figure 2B-C; a 2-way MANOVA; a main effect of Counterconditioning [F(2,16)=8.6, p<0.01]; no main effect of Retrieval [F(2,16)=1.25, p>0.05]; a Retrieval X Counterconditioning interaction [F(2,16)=3.79, p<0.05]. Post-hoc for the Counterconditioning main effect: *Zif268* [no CC vs CC, p<0.01]; *Bdnf* [no CC vs CC, p<0.05]. Post-hoc analysis for Retrieval X Counterconditioning effect: *Bdnf* [No Retrieval+CC vs Retrieval+CC, p<0.05]). There was no change in the expression of *Zif268* or *Bdnf* in the dorsal hippocampus (Figure 2D-E; a 2-way MANOVA; no main effect of Counterconditioning [F(2,17)=3.21, p>0.05], or Retrieval [F(2,17)=0.19, p>0.05]; no Retrieval X Counterconditioning interaction [F(2,17)=0.41, p>0.05]). These results suggest that the increase of the expression of *Bdnf* in the mPFC provides a molecular correlate for the combined behavioral effect of retrieval-counterconditioning, i.e., for memory updating.

### Experiment 3: Aversive counterconditioning following memory retrieval suppresses renewal of alcohol seeking in an operant self-administration procedure

Having shown that aversive counterconditioning applied after the retrieval of alcohol-associated memories prevented the reinstatement of alcohol memories in a place conditioning paradigm, we next tested whether a similar effect can be obtained in an instrumental learning paradigm that models consummatory behaviors, i.e., in operant alcohol self-administration.

To this end, we developed a two-phases operant counterconditioning procedure, by adopting an ABA renewal procedure (Bouton and Bolles, 1979; Crombag et al., 2008), widely used to explore context-induced reinstatement of drug seeking (Burattini et al., 2006; Janak and Chaudhri, 2010; Marchant et al., 2013b; Goltseker et al., 2019). First, rats were trained to self-administer alcohol in context A, by responding on an active lever to obtain alcohol rewards (Figure 3A, experimental design). We found that by the end of the acquisition phase rats established stable responding on the lever reinforced with alcohol, as confirmed by the analysis of the active and inactive lever-presses during the last 6 training sessions under a FR3 ratio (Figure 3B&C; a 2-way mixed model ANOVA, main effects of Lever [F(5,255)=161.55, p<0.0001] and Day [F(5,255)=4.45; p<0.01], and a Lever X Day interaction [F(5,255)=4.01, p<0.01]).

**Figure 3.**
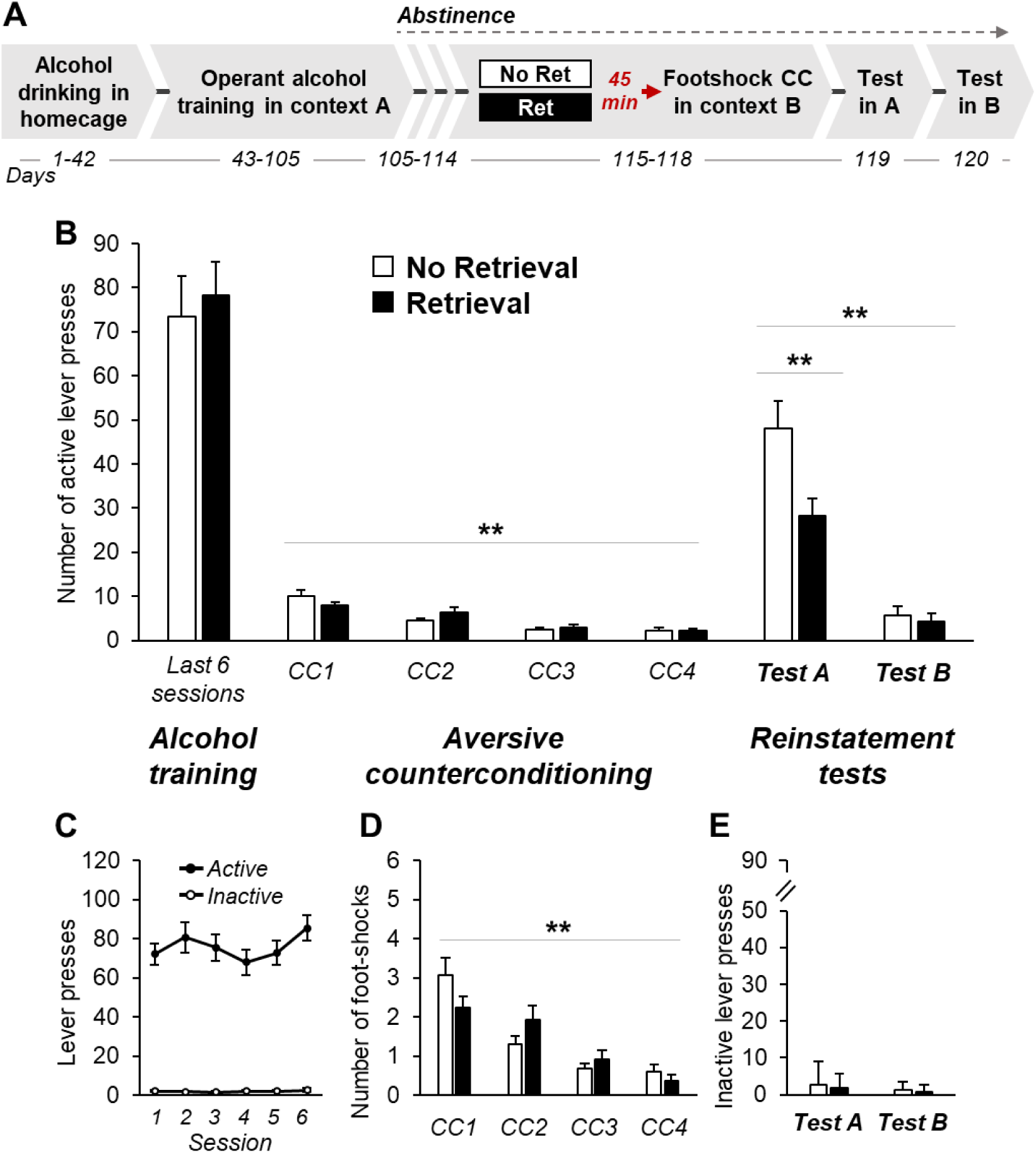
Aversive counterconditioning performed 45 min after alcohol-memory retrieval reduces reinstatement of operant alcohol seeking in rats. **A.** Schematic representation of the experimental procedure (CC, counterconditioning; Ret, retrieval). Following prolonged alcohol drinking and operant self-administration, lever-pressing was counterconditioned with foot-shocks in context B. Forty-five min before each counterconditioning session, rats were briefly exposed to the alcohol odor-taste cues (Retrieval group), or water (No Retrieval group) in the homecage. Next, reinstatement of alcohol seeking was assessed in context A, followed by a test session in context B. **B**. Number of active lever presses during the alcohol training in context A, counterconditioning in context B, test in context A, and test in context B. **C**. Number of active and inactive lever presses during the last 6 sessions of alcohol training in context A. **D**. Number of foot-shocks delivered during the counterconditioning phase in context B. **E**. Number of inactive lever-presses during tests in context A and B. Data are expressed as mean +S.E.M; *p<0.05; **p<0.01; n=13 per group.

After 10 days of abstinence from alcohol, rats were assigned to 2 groups (Retrieval, No Retrieval). Memory retrieval consisted of a brief exposure to the alcohol odor-taste cues in the home cage by presenting a bottle with the tip covered with alcohol (Retrieval group), or water (No Retrieval group; see Methods). We previously demonstrated that this manipulation reactivated alcohol-associated memories (Barak et al., 2013). Forty-five min later, rats were placed in a context distinct from the training context by olfactory and visual features. In this distinct environment (context B), rats received aversive counterconditioning training, in which every 3rd active-lever press resulted in the delivery of a mild foot-shock (limited to max of 6 foot-shocks per a 30-min session). During the four counterconditioning sessions in Context B, rats gradually ceased to respond on the active lever (Figure 3B; a 2-way mixed model ANOVA: a main effect of Day [F(3,72)=35.74, p<0.0001], a Day X Group interaction [F(3,72)=2.77, p<0.05], but no main effect of Group [F(3,72)=0.01, p>0.05]). Importantly, during the counterconditioning phase, rats from both groups received a similar number of foot-shocks in total (Figure 3D; Retrieval vs No Retrieval [t(25)=0.2, p>0.05]).

Relapse to alcohol-seeking responding was then assessed in alcohol-primed test sessions performed in the alcohol-paired context A and in the aversion-associated context B. We found that during a 30-min session of non-reinforced lever responding in context A, rats that received aversive counterconditioning following alcohol-memory retrieval showed fewer active lever-presses compared to control rats that received aversive counterconditioning without memory retrieval, whereas in context B responding was similar in both groups. Also, the overall active lever-pressing was higher in context A (Figure 3B; a 2-way mixed model ANOVA: main effects of Group [F(1,24)=6.38, p<0.05] and Test [F(1,24)=85.74, p<0.0001], and a Test X Group interaction [F(1,24)=6.83, p<0.05]. Post-hoc analysis: No Retrieval vs. Retrieval groups [Test A, p<0.01], [Test B, p>0.05]). No effects were found on the inactive lever presses (Figure 3E; all p’s>0.05).

To conclude, we found that the renewal of alcohol seeking in context A was significantly decreased when aversive counterconditioning in context B was given shortly after alcohol memory retrieval, i.e., within the “reconsolidation window”.

### Experiment 4: Aversive counterconditioning outside the “reconsolidation window” does not prevent renewal of alcohol seeking in an operant self-administration procedure

Next, we tested whether a similar effect can be obtained when aversive counterconditioning is given 6 hours after alcohol-memory retrieval, i.e., outside the “reconsolidation window”.

To this end, we trained rats as in Experiment 3, but memory retrieval was given 6 hours before the aversive counterconditioning sessions (Figure 4A, experimental design).

**Figure 4.**
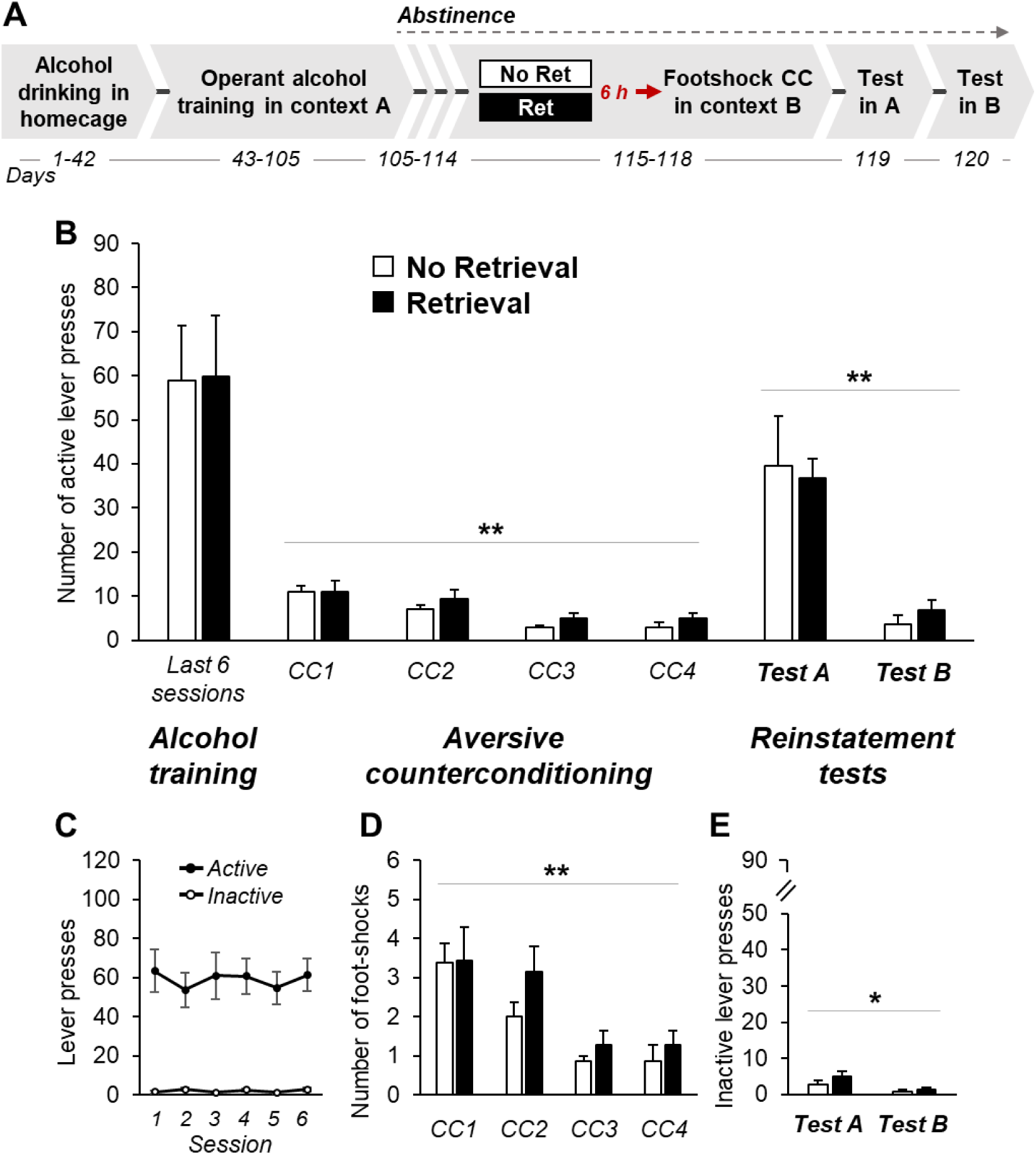
Aversive counterconditioning performed 6 h after alcohol-memory retrieval failed to prevent reinstatement of operant alcohol seeking in rats. **A.** Schematic representation of the experimental procedure (CC, counterconditioning; Ret, retrieval). Following prolonged alcohol drinking and operant self-administration, lever-pressing was counterconditioned with foot-shocks in context B. Six hours before each counterconditioning session, rats were briefly exposed to the alcohol odor-taste cues (Retrieval group), or water (No Retrieval group) in the homecage. Next, reinstatement of alcohol seeking was assessed in context A, followed by a test session in context B. **B**. Number of active lever presses during the alcohol training in context A, counterconditioning in context B, test in context A, and test in context B. **C**. Number of active and inactive lever presses during the last 6 sessions of alcohol training in context A. **D**. Number of foot-shocks delivered during the counterconditioning phase in context B. **E**. Number of inactive lever-presses during tests in context A and B. Data are expressed as mean +S.E.M; *p<0.05; **<0.01; n=13 per group.

After 9 weeks of training to self-administer alcohol in context A, rats established stable responding on the lever reinforced with alcohol, as confirmed by the analysis of the active and inactive lever-presses during the last 6 training sessions (Figure 4B&C; a 2-way mixed model ANOVA, a main effect of Lever [F(5,140)=40.32, p<0.0001], but no main effect of Day [F(5,140)=1.11; p>0.05], and no Lever X Day interaction [F(1,140)=1.22, p>0.05]).

During the four counterconditioning sessions in Context B, rats gradually ceased to respond on the active lever (Figure 4B; a 2-way mixed model ANOVA: a main effect of Day [F(3,39)=15.33, p<0.0001], but no main effect of Group [F(3,39)=1.19, p>0.05], and no Day X Group interaction [F(3,39)=0.41, p>0.05]). Also, during the counterconditioning phase, rats from both groups received a similar number of foot-shocks in total (Figure 4D; Rerieval vs No Retrieval: [t(13)=1.19, p>0.05]).

Relapse to alcohol-seeking responding was then assessed in the alcohol-paired context A and compared to responding in the aversion-associated context B. We found that non-reinforced lever-responding did not differ between rats that received alcohol-memory retrieval 6 hours before aversive counterconditioning and control rats, who underwent aversive counterconditioning without prior memory retrieval. Furthermore, we found that the overall active lever-pressing was higher in context A compared to context B, in which lever-responding was also similar in both groups (Figure 4B; a 2-way mixed model ANOVA: a main effect of Test [F(1,13)=24.01, p<0.0001], no main effect of Group [F(1,13)=0.01, p>0.05], and no Test X Group interaction [F(1,13)=0.21, p>0.05]). No effects were found on the inactive lever presses (Figure 4E; all p’s>0.05).

To conclude, we found that alcohol seeking was equally reinstated in both groups, suggesting that alcohol-memory retrieval given 6 hours before aversive counterconditioning did not affect the recovery of alcohol seeking.

## Discussion

We show here that relapse to alcohol seeking can be prevented when aversive counterconditioning is delivered during the reconsolidation of alcohol memories. Specifically, in the CPP paradigm, which is based on classical conditioning and does not include voluntary alcohol consumption, mice did not show reinstatement of alcohol seeking when aversive counterconditioning in the alcohol-paired context was preceded by alcohol-memory retrieval. Similarly, in the operant alcohol self-administration paradigm, in which rats voluntarily consume alcohol, rats trained to lever-press for alcohol showed decreased renewal of alcohol seeking when the lever-pressing was counterconditioned with mild foot-shocks, shortly but not long after memory retrieval. Together, our results indicate that aversive counterconditioning can suppress relapse to alcohol seeking, only when performed following memory retrieval, i.e., within the “reconsolidation window”. Furthermore, our results suggest that increased *Bdnf* expression in the mPFC following retrieval-counterconditioning may provide a molecular correlate for the memory updating/replacement effect.

### Behavioral interference with alcohol memory reconsolidation prevents relapse

Our finding that reinstatement of the alcohol-CPP can be suppressed by preceding aversive counterconditioning with alcohol-memory retrieval is consistent with our previous report, in which aversive counterconditioning applied shortly after cocaine-memory retrieval prevented reinstatement of the cocaine-CPP (Goltseker et al., 2017). We also show, as previously (Goltseker et al., 2017), that counterconditioning with no prior memory retrieval does not prevent drug-seeking reinstatement. Indeed, a regular counterconditioning procedure is known to be only temporary effective in suppressing the target behavior (Bouton and Peck, 1992; Brooks et al., 1995; van Dis et al., 2019), suggesting that the original memory remains intact and can recover (Bouton and Peck, 1992; Brooks et al., 1995). On contrary, as our findings suggest, in the retrieval-counterconditioning paradigm, aversive information presented while the drug-memory is in an active state can be integrated into the original memory, thus updating or replacing it, and consequently preventing its behavioral expression (Das et al., 2015; Goltseker et al., 2017; Lee et al., 2017; Gisquet-Verrier and Riccio, 2018). Relatedly, we show that post-retrieval counterconditioning leads to a switch in behavior: from alcohol-CPP to CPA. This finding further supports our suggestion that the appetitive alcohol memory was updated, or replaced by an aversive memory, and that the valance of the cue has been changed.

Importantly, addiction-related responses are governed not only by maladaptive Pavlovian conditioning-based memories, but also by instrumental conditioning mechanisms that form persistent habits (Everitt et al., 2001; Hyman, 2005; Corbit and Janak, 2007; Everitt and Robbins, 2016). Targeting the memories underlying operant behaviors via reconsolidation mechanisms has been particularly challenging (Hernandez and Kelley, 2004; Exton-McGuinness et al., 2015; Vousden and Milton, 2017; Exton-McGuinness and Milton, 2018; Exton-McGuinness et al., 2019). Indirectly, it is possible to attenuate operant behaviors by disrupting the memories for the conditioned stimuli, in the presence of which the target operant behaviors are suppressed or facilitated (Hellemans et al., 2006; Lee et al., 2006; Milton et al., 2008b; Milton et al., 2008a; Milton et al., 2012; Dunbar and Taylor, 2017b). Strikingly, the integral memory components underpinning operant responding were also shown to be prone to reconsolidation-dependent modification (Exton-McGuinness et al., 2014; Exton-McGuinness and Lee, 2015; Exton-McGuinness et al., 2019). Thus, attenuation of operant responding can be achieved via pharmacological interference with memory reconsolidation (Lee et al., 2006; Milton et al., 2008b; Milton et al., 2008a; Milton et al., 2012; Barak et al., 2013; Exton-McGuinness et al., 2014; Exton-McGuinness and Lee, 2015; Dunbar and Taylor, 2017a), with only a few studies testing behavioral manipulations, such as retrieval-extinction (Xue et al., 2012; Millan et al., 2013; Luo et al., 2015). Here, we show that the retrieval-counterconditioning manipulation attenuates the context-induced reinstatement (renewal) of operant alcohol self-administration, even after long-term operant training. Thus, our findings further support the notion that operant behaviors can be attenuated via behavioral interference with memory reconsolidation.

This finding has a critical translational value, as unlike the CPP paradigm, in which exposure to a drug of abuse is short and involuntary (Tzschentke, 2007), operant self-administration procedures are used to model a variety of aspects in addiction (Shaham et al., 2003; Crombag et al., 2008; Everitt et al., 2008; Marchant et al., 2013a; Venniro et al., 2016), including alcohol relapse after prolonged voluntary consumption (Koob, 2000; Burattini et al., 2006; Epstein et al., 2006; Goltseker et al., 2019). Moreover, the retrieval-counterconditioning paradigm is feasible in humans, as it was recently shown that re-association of the alcohol cues with aversion upon alcohol memory reactivation, yielded reduction in alcohol craving in hazardous drinkers (Das et al., 2015; Das et al., 2018).

One of the major challenges in relapse prevention derives from the fact that psychotherapy is typically given in the clinics, which is a different environmental context than the natural alcohol-/drug-taking environment. As a result, most patients relapse when they return home, or to their usual drug-taking environment, even after showing successful decreases in craving at the clinics (Carter and Tiffany, 1999; Witteman et al., 2015). We show here that the reconsolidation-based relapse prevention treatment could be effective also when applied in a different context than the original alcohol-taking context. Specifically, rats were trained in a modified ABA-renewal procedure (Bouton and Bolles, 1979; Crombag et al., 2008), in which they self-administered alcohol in context A, and then received the “behavioral treatment”, aversive counterconditioning, in a distinct context (context B). As expected, we found that without memory retrieval, rats reinstated alcohol seeking upon their return to the alcohol-associated environment (context A), similar to relapse in patients. However, context-induced relapse to alcohol seeking (renewal) was considerably reduced when the counterconditioning treatment was preceded by alcohol memory retrieval, and was conducted within the “reconsolidation window” (shortly, but not long-after, alcohol-memory retrieval). Importantly, both the memory retrieval and counterconditioning sessions were given outside the alcohol-taking context A. Thus, our procedure addresses the clinical issue of context-dependent relapse.

Interestingly, unlike in the alcohol-CPP paradigm, in the operant self-administration paradigm we did not observe full disruption of alcohol seeking. Similar reduction in relapse but not its complete prevention was also observed in previous studies, in which operant drug seeking was targeted with retrieval-extinction manipulations (Xue et al., 2012; Millan et al., 2013; Luo et al., 2015). This finding may suggest that in the operant procedure, the retrieval-counterconditioning manipulation did not fully overwrite the memories underpinning operant alcohol seeking but rather updated them.

The finding that the reduction in relapse to alcohol seeking in the present operant procedure was partial, raises the possibility that a brief exposure to the alcohol odor-taste cue did not reactivate all the memory elements that underlie operant alcohol seeking, thus preventing an exhaustive memory update. However, this possibility is challenged by our previous findings, showing that a similar memory retrieval procedure, followed by administration of rapamycin, an inhibitor of the mechanistic target of rapamycin complex 1 (mTORC1), led to a complete abolition of alcohol seeking in a similar retention test (Barak et al., 2013). Moreover, alcohol’s taste and odor are intrinsic components of oral alcohol consumption, that reliably predict the onset of the reinforcing effect of alcohol (Katner and Weiss, 1999; Backstrom et al., 2004; Filbey et al., 2008; Barak et al., 2013; Brasser et al., 2015), and elicit stronger responses compared to other alcohol-associated cues (Katner et al., 1999). Therefore, it is likely that exposure to the odor-taste cue triggered the reactivation of a wide range of alcohol-related memories, including those underpinning operant alcohol seeking.

If so, it is possible that aversive counterconditioning lever-pressing with footshocks did not swap completely the valence of the alcohol memories, that control the operant alcohol seeking. Here, we re-associated operant responding with footshocks, a well-established method to counter-condition or punish appetitive behaviors (Bouton and Peck, 1992; Marchant et al., 2013b; Bali and Jaggi, 2015). However, here we used very mild footshocks (0.25 mA), considered to be just above the responsivity threshold intensity (Gravius et al., 2006), and capable to elicit only moderate avoidant response in rats (Muravieva and Alberini, 2010). On the contrary, in the previous counterconditioning studies performed on operant paradigms the foot-shock intensity was higher (0.5 mA) (Williams and Barry III, 1966; Brooks et al., 1995). We chose a low footshock intensity to prevent a robust freezing response, which could suppress completely further instrumental behavior (D’amato and Fazzaro, 1966; Rushen, 1986), thus averting the counterconditioning of lever-pressing. Also, we sought to avoid the unnecessary pain, which could affect the rewarding and learning systems balance (Brischoux et al., 2009), thus confounding the reactivation-counterconditioning effect. Yet, it is possible that a stronger counterconditioning (e.g., higher footshock intensity) given within the “reconsolidation window”, would result in full abolishment of alcohol-seeking reinstatement.

### *Bdnf* expression in the mPFC is a possible molecular correlate for the effects of the retrieval-counterconditioning manipulation

We found that aversive counterconditioning led to upregulation of *Bdnf* mRNA levels in the mPFC in mice, only when it was preceded by alcohol memory retrieval. Interestingly, *Bdnf* expression has been previously implicated in new memory formation (Berglind et al., 2007; Choi et al., 2012; Baker-Andresen et al., 2013), but not in memory reconsolidation (Lee et al., 2004). However, here we found no change in *Bdnf* expression after counterconditioning *per se* (no retrieval group), which is thought to form a new memory trace that competes with the original memory for behavioral expression (Bouton and Peck, 1992; Brooks et al., 1995). On the contrary, we found increased *Bdnf* expression when counterconditioning was performed shortly after memory retrieval, which is thought to allow integration of new information into the existing memory, updating, or even replacing it (Das et al., 2015; Goltseker et al., 2017; Lee et al., 2017; Gisquet-Verrier and Riccio, 2018; Goltseker et al., 2019). Thus, our findings suggest that the increase in *Bdnf* expression in the mPFC following retrieval-counterconditioning correlates with memory updating, rather than with the formation of a new (competing) memory.

It was previously reported that activation of BDNF signaling, by a systemic agonist of the BDNF receptor TrkB, potentiated the effect of the retrieval-extinction procedure and abolished the recovery of fear responses mice (Baker-Andresen et al., 2013). Likewise, direct infusion of BDNF into the mPFC reduced the expression of the previously established fear response in rats (Peters et al., 2010), possibly by partial reversal of the neuronal changes induced during fear conditioning (Peters et al., 2010). In our study, upregulation of the endogenous *Bdnf* expression in the mPFC after retrieval-counterconditioning correlated with the suppression of alcohol seeking, suggesting that BDNF may have played a similar role in reversing alcohol CPP-induced changes. Nevertheless, the causal role of the BDNF in the mPFC in the effect of retrieval-counterconditioning procedure is yet to be empirically established.

The expression levels of *Bdnf* and *Zif268* in the mPFC and dorsal hippocampus were not affected by the retrieval of alcohol memories, contrary to the previous reports about the involvement of these genes in memory reconsolidation (Bozon et al., 2003; Lee et al., 2004; Tedesco et al., 2014; Radiske et al., 2015; Radiske et al., 2017). It is very likely that we did not observe this effect, because in our study the levels of the *Bdnf* and *Zif268* mRNA were assessed 90 min after alcohol memory retrieval (since mice were also counterconditioned). Given the rapid expression dynamics of these genes in memory tasks (Tischmeyer and Grimm, 1999; Hall et al., 2000; Ressler et al., 2002; Barry and Commins, 2017), it is possible that this time-point, chosen to reflect the transcription dynamics of counterconditioning, was beyond the peak of expression of *Bdnf* and/or *Zif268* mRNAs caused by retrieval.

To conclude, our current results provide a critical expansion of the retrieval-counterconditioning paradigm, by showing that: 1. it can be applied on additional drugs of abuse (alcohol); 2. It can be used in different and more complex behavioral paradigms, which are more relevant for translational research in addiction (i.e., operant alcohol self-administration)3. We can identify molecular correlates for the behavioral effect of aversive counterconditioning occurring after memory retrieval. Finally, we demonstrate the effectiveness of this behavioral manipulation in reducing the return of alcohol seeking, even when performed away from the alcohol context. Therefore, we introduce a potential therapeutically-relevant strategy to address a critical clinical issue of context-induced relapse.

## Conflict of Interest

The authors declare no competing financial interests.

## Acknowledgement

The research described in and the writing of this review were supported by funds from the Israel Science Foundation (ISF) grants 968-13 and 1916-13 (S.B.).

## Author contribution

KG, HH, and SB designed the research, KG and HH conducted the experiments. KG and SB analyzed the data and wrote the manuscript.

